# Effects of the Ayurvedic Formulation Swarna Sindhura on Serum Gonadotropins and Testosterone Levels in Male Sprague-Dawley Rats

**DOI:** 10.1101/2025.09.06.674625

**Authors:** Maharun Nessa, Israt Jahan, Musammat Umme Habiba, Shahabuddin Kabir Choudhuri

**Affiliations:** Department of Pharmacy, Jahangirnagar University, Savar, Dhaka-1342

**Author notes:** **Corresponding Author: Shahabuddin Kabir Choudhuri**, Former Professor, Department of Pharmacy, Jahangirnagar University, Savar, Dhaka-1342. **Competing interests** The authors declare no competing interests.

**Keywords:** Swarna Sindhura (SRNm), Ayurvedic medicine, Gonadotropin hormone, Testosterone, Progesterone, Luteinizing hormone (LH), Follicle-stimulating hormone (FSH), 17-beta-Estradiol (E2), Dehydroepiandrosterone sulfate (DHEA-S), Sprague-Dawley Rats

## Abstract

Swarna Sindhura (SRNm) is a traditional Ayurvedic formulation used in the management of hypertension, particularly in rural populations in the Indian subcontinent. Despite its extensive use, the endocrine effects of SRNm still remain unknown due to a lack of studies. This study examined the effect of SRNm administration on steroid and gonadotropin hormone profiles in male Sprague-Dawley rats. Animals received oral SRNm at a dose of 40 mg/kg daily for 28 days, whereas control rats received normal food and water. Serum concentrations of dehydroepiandrosterone sulfate (DHEA-S), total testosterone, progesterone, 17β-estradiol (E2), luteinizing hormone (LH), and follicle-stimulating hormone (FSH) were quantified and statistically analyzed to understand the effects of SRNm. SRNm treatment elicited a statistically significant elevation in serum progesterone, suggesting enhanced steroidogenic activity potentially relevant to its traditional antihypertensive use. DHEA-S and testosterone are slightly decreased, indicating a subtle modulatory influence on androgen biosynthesis without compromising basal production. Gonadotropin analysis revealed that LH levels have decreased, while FSH levels have slightly increased, but the change is non-significant. These findings show that SRNm had a selective effect on progesterone synthesis, while maintaining overall hormonal homeostasis. Further detailed studies are essentially needed to clarify its endocrine effects and therapeutic potential.

## Introduction

Ayurveda, traditional medicine in Indian subcontinent, signifies one of the primogenital holistic healing practices in the world. Engrained in the Vedic tradition, it is projected to have developed over several thousand years. It has since changed into a wide-ranging system of healthcare with both theoretical and practical dimensions based on religious beliefs, philosophy, and the natural sciences [1]. Its therapeutic approaches include lifestyle and dietary interventions alongside a wide range of herbal formulations. Today, Ayurveda is officially recognized in India and practiced globally, with increasing interest in its formulations for both therapeutic use and drug discovery[2].

In Bangladesh, traditional medicine occupies a significant role within the national healthcare framework. The production, distribution, and regulation of these medicines are governed by key legislations such as the Drug Act (1940), the Drug Regulation (1946), and the Drug (Control) Amendment Act (2006). To further strengthen oversight, the Government of Bangladesh established the Office of the Director of Homeo and Traditional Medicine in 1991 under the Directorate General of Health Services (Pathak, 2022)[3]. This reflects a broader regional acknowledgement of the value of indigenous medical systems, which continue to coexist alongside modern biomedicine in the South Asian healthcare system.

Among Ayurvedic medicines, Swarna Sindhura (SRNm) is widely considered for its traditional use in male reproductive health, general incapacity, and skinniness. SRNm is prepared through traditional processes combining herbal and mineral ingredients, including purified metals such as gold and mercury. While historical emphasis has been placed on the therapeutic benefits of this formulation, its potent composition permits careful use under strict medical supervision. The safety of heavy metal-containing preparations has always been a serious concern that needs to be further highlighted, with importance for rigorous preclinical evaluation [4, 5].

Reproductive health and hormonal balance in males are influenced by gonadotropins and testosterone, which are critical for spermatogenesis and overall fertility. Despite the traditional use of Swarna Sindhura in improving male reproductive system, there is very limited scientific evidence regarding its effects on serum gonadotropins and testosterone levels. Preclinical studies using animal models provide an important agenda to evaluate both the safety and efficacy of these formulations under controlled conditions [6, 7].

The present study, therefore, aims to investigate the effects of the Ayurvedic formulation Swarna Sindhura on serum gonadotropins (LH and FSH) and testosterone levels in male Sprague-Dawley rats. This research not only seeks to validate the traditional claims of Swarna Sindhura in enhancing male reproductive health but also aims to provide a scientific basis for its safe use and further clinical exploration.

## Materials and Methods

### Collection and Preparation of Test Substance

The Ayurvedic formulation Swarna Sindhura (SRNm) was procured from the licensed manufacturer Sri Kundeswari Aushadhalaya Ltd. in Chittagong, Bangladesh. The formulation was prepared in accordance with traditional Ayurvedic protocols as described in standard pharmacopeias [8-11]. Its composition consists of Shuddha Parada, which is purified and processed mercury, at 48 grams; Shuddha Gandhaka, which is purified and processed sulphur, at 48 grams; and Swarna, which is gold bhasma, at 12 grams. These primary ingredients are triturated with Vata praroha rasa, the juice extract of Ficus benghalensis, and Kumari rasa, the juice extract of Aloe vera, using a quantity sufficient for the process. The mixture is then prepared using the special Kupipakwa technique, a specific Ayurvedic method of processing herbo-mineral preparations in a glass bottle under controlled heat. For the purposes of oral administration in this study, a suspension was prepared by grinding the tablets and following the procedure outlined in the Bangladesh National Ayurvedic Formulary (1992).

### Experimental Animals and Ethical Considerations

Healthy male Albino rats (Rattus norvegicus, Sprague-Dawley strain), eight weeks old and weighing 50–70 g, were used for this study. The animals were housed in the Animal House of the Department of Pharmacy, Jahangirnagar University, Savar, Dhaka, Bangladesh under approved animal protocol. They were maintained in plastic cages (30 x 20 x 13 cm) with soft wood shaving bedding under a natural light/dark cycle, a well-ventilated environment, and ad libitum access to food (mouse chow was prepared according to the formula established at BCSIR, Dhaka) and water. All experimental procedures were conducted in institutional compliance with the ethical guidelines for the care and use of laboratory animals. The study protocol was approved by the Institutional Animal Ethics Committee.

### Experimental Design and Dosing

Rats were randomized and allocated into two groups of ten animals each: a control (n=10), which received an oral administration of distilled water (placebo), and a treated (n=10), which received an oral administration of Swarna Sindhura suspension at a dose of 40 mg/kg body weight. The dose volume was calibrated to ensure precise dosing without significantly contributing to the total body fluid volume [12]. Administration was performed daily using an intragastric syringe between the hours of 10:00 AM and 12:00 PM for 28 consecutive days [13, 14]. To enable accurate longitudinal monitoring, individual rats were identified by unique tail markings and tagging.

### Sample Collection

After the 28-day treatment period, animals were fast for 18 hours. Twenty-four (24) hours after the final administration, rats were anesthetized with ketamine (500 mg/kg, intraperitoneally) [15-17]. Whole blood samples were collected from the post-vena cava, transferred immediately to sample tubes, and allowed to clot. The serum was separated using a dry Pasteur pipette, stored refrigerated, and all analyses were completed within 24 hours of collection [18].

### Biochemical Analysis

Serum levels of Dehydroepiandrosterone Sulfate (DHEA-S), Testosterone, Progesterone, Estradiol, Luteinizing Hormone (LH), and Follicle Stimulating Hormone (FSH) were determined using the Chemiluminescent Microparticle Immunoassay (CMIA) method from Abbott Laboratories (USA)[19].

### Statistical Analysis

All group data are presented as Mean ± Standard Error of the Mean (SEM). The statistical comparisons between the control and treated groups in rats were made using an unpaired student’s t-test. Data analysis was conducted using SPSS for Windows (Version 11). Differences were considered statistically significant at p < 0.05, with p < 0.01 and p < 0.001 considered highly and very highly significant, respectively.

## Results

This study evaluated the impact of chronic administration (28 days) of the Ayurvedic herbo-mineral formulation Swarna Sindhura (SRNm) on the steroidal and gonadotropin hormone profiles in male Sprague-Dawley rats.

### Steroid Hormone Profile

Chronic oral administration of SRNm at a dose of 40 mg/kg elicited distinct modulatory effects on the steroid hormone profile, with the most pronounced alteration observed in serum progesterone levels.

### Dehydroepiandrosterone Sulfate (DHEA-S) and Testosterone

SRNm administration resulted in a modest, statistically non-significant reduction in serum DHEA-S and total testosterone compared to controls (**Fig. 1**). Given the role of DHEA-S as a precursor in androgen biosynthesis, this coordinated downward trend though not statistically significant, suggests an elusive regulatory influence on the adrenal and gonadal androgenic pathways. The absence of significant changes shows that SRNm does not considerably impair the basal synthesis of these vital androgens.

**Figure 1.**
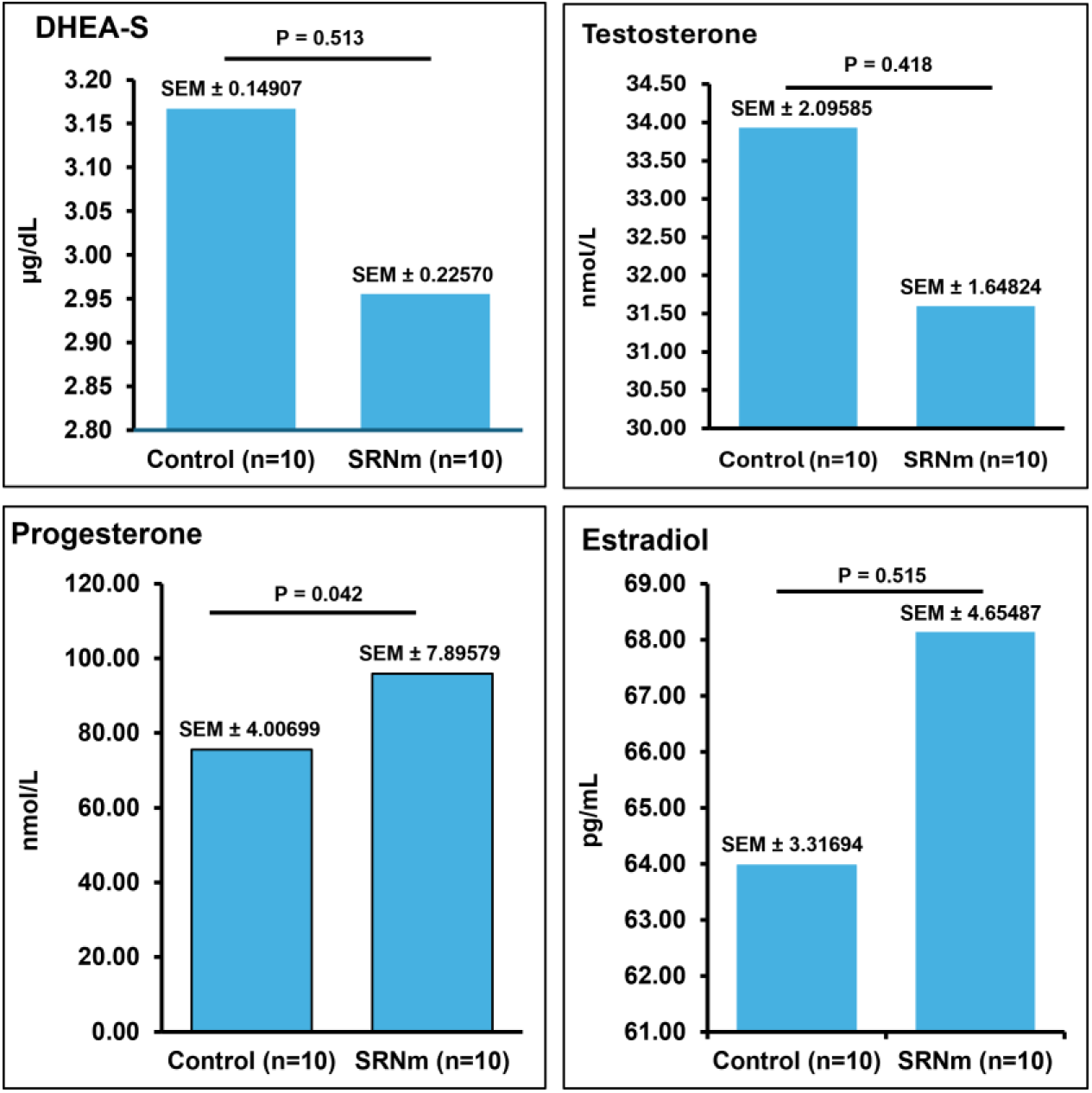
Effect of Swarna Sindura (SRNm) on serum steroid hormone profile in male rats. Serum levels of DHEA-S, testosterone, progesterone, and estradiol were measured in male Sprague–Dawley rats following chronic administration of SRNm.

### Progesterone

A significant increase in serum progesterone was observed following SRNm treatment (**Fig. 1**). In males, progesterone is synthesized in both the adrenal cortex and testes, contributing to gonadotropin regulation and steroidogenic activity. The observed increase may suggest increased progesterone biosynthesis or a shift in steroidogenic enzyme activity. Given progesterone’s role in controlling vascular tone and fluid balance, this finding may have effects for SRNm’s traditional use in managing hypertension and deserves further mechanistic exploration at the molecular level.

### Estradiol (E2)

Serum estradiol levels showed an insignificant increase in the SRNm-treated group compared to control rats (**Fig. 1**). This negligible change, in the context of stable testosterone levels, suggests that the androgen-to-estrogen ratio persisted largely unaltered. The increase may be secondary to changes in precursor dynamics activity, but does not appear to be a primary pharmacological effect of SRNm.

### Gonadotropin Hormone Profile

Evaluation of pituitary gonadotropins showed minimal impact of SRNm on the hypothalamic-pituitary-gonadal (HPG) axis, with no significant changes in Luteinizing Hormone (LH) or Follicle-Stimulating Hormone (FSH).

### Luteinizing Hormone (LH)

SRNm administration led to a 26.03% non-significant reduction in serum LH levels (**Fig. 2**). As LH is necessary for Leydig cell-mediated testosterone production, the lack of significant changes suggests that Leydig cell function and integral upstream regulatory mechanisms are conserved.

**Figure 2.**
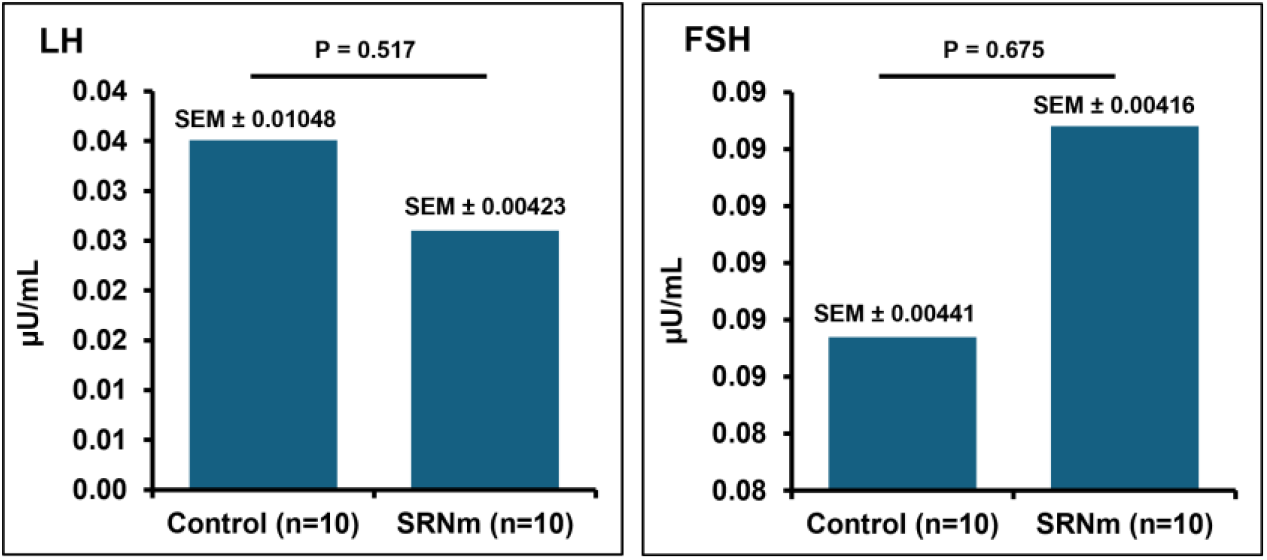
Effect of Swarna Sindura (SRNm) on serum Gonadotropin hormone profile in male rats. Serum levels of Luteinizing Hormone (LH), and Follicle-Stimulating Hormone (FSH) were measured in male Sprague–Dawley rats following chronic administration of SRNm.

### Follicle-Stimulating Hormone (FSH)

FSH levels remained steady, with a negligible increase in the treatment group compared to control rats (**Fig. 2**). FSH plays an important role in Sertoli cell function and spermatogenesis. The insignificant changes result in SRNm not unfavorably affecting testicular support functions or spermatogenic regulation.

## Discussion

Ayurvedic medicine has gained immense importance as an alternative to conventional medical treatment. Improving their safety and efficacy requires more careful consideration of their historical applications alongside modern scientific toxicological investigations [20, 21]. This study employed hormonal screening to assess the safety and efficacy profile of the herbo-mineral formulation Swarna Sindhura (SRNm) in rats, in line with approaches recommended for evaluating traditional medicines [22, 23].

The most significant finding of this study is the statistically significant increase in serum progesterone levels following chronic SRNm administration. While progesterone is mainly well-known for its role in the female reproductive cycle, where it is vital for preparing the uterus for implantation, maintaining pregnancy, and serving as a strategic marker of ovulation and luteal phase deficiency [24], its role in males is also critical. In males, progesterone is produced by the adrenal cortex and the testes, acting as an essential neurosteroid and precursor for other steroid hormones [25, 26].

This distinct increase in progesterone offers a decisive understanding of SRNm’s mechanism and functional activities. The significant change suggests that a targeted effect on the enzymes or biosynthetic pathways specific to progesterone production, rather than a generalized endocrine disruption, will steady with the adaptogenic properties [27, 28]. The non-significant changes observed in testosterone, DHEA-S, estradiol, LH, and FSH levels further demonstrate that SRNm does not suppress the broader hypothalamic-pituitary-gonadal (HPG) axis and conserves inclusive endocrine homeostasis, supporting its safety profile as a traditional medicine [29, 30].

The associations of increased progesterone are complicated to understand without proper functional studies. As a neuro-steroid, progesterone and its metabolites potentiate GABAergic neurotransmission, which can endorse vasodilation and reduce sympathetic tone [31, 32]. This mechanism proposes a reasonable explanation for the traditional use of SRNm in managing hypertension, suggesting its benefits may be facilitated through this endocrine-neurological pathway, similar to other cardiovascular-active herbal preparations [33-35]. Furthermore, the stability of the HPG axis, evidenced by unchanged LH and FSH levels, indicates a favorable safety profile regarding reproductive hormone regulation, an important consideration for long-term use of traditional medicines (36, 37).

## Conclusion

Swarna Sindhura demonstrates a significant modulatory effect on steroidogenesis, especially by increasing progesterone. This effect, together with the absence of significant adverse effects on other hormonal axes, supports its potential as a targeted therapeutic agent with controlled preclinical studies. These findings highlight the importance of scientific validation for traditional Ayurvedic medicines, connecting historical use with evidence-based application. Additional research is necessary to association this progesterone modulation to cardiovascular endpoints and to reveal the precise molecular mechanisms involved, particularly through detailed investigation of steroidogenic enzyme expression and activity.

## Author contributions

MN, and SKC participated in the research design. MN conducted the experiments, and MN, IJ and MUH performed data analysis and write the manuscript. SKC supervised the study and edited the paper. All authors read and approved the paper.

